# Interact: Automated analysis of protein-ligand interactions by 1D and 2D NMR

**DOI:** 10.1101/215194

**Authors:** Pierre Millard, Guy Lippens

**Affiliations:** LISBP, Université de Toulouse, CNRS, INRA, INSA, Toulouse, France

**Keywords:** Protein interaction, Regulation, Structural biology, Systems biology, Biochemistry

## Abstract

NMR titration experiments contain rich information on the thermodynamic, kinetic and structural aspects of protein-ligand interactions. Automated tools are required to process the large number of signals typically acquired in these experiments and facilitate quantitative interpretations. We present Interact, a Python script accessible within the Bruker BioSpin TopSpin™ software, which allows automated analysis of both 1D and 2D NMR titration experiments. Interact performs peak picking and annotation of the successive spectra and supports quantitative interpretation of changes in chemical shifts and linewidths induced by the ligand (e.g. to estimate dissociation constants) through different fitting procedures. Interact can be applied to all types of 1D and 2D NMR experiments and all nuclei, hence facilitating routine analysis of existing and forthcoming NMR titration data. Interact was implemented in Python and can be used on Windows, Unix and MacOS platforms. The source code is distributed under OpenSource license at http://github.com/MetaSys-LISBP/Interact.

## Introduction

NMR is a powerful tool to study thermodynamic, structural, and kinetic aspects of protein-ligand (Fielding, 2003; Furukawa *et al*., 2016) or protein-protein (Takeuchi and Wagner, 2006; Teilum *et al*., 2017) interactions. In titration experiments, meaningful information is typically extracted from the changes of chemical shift or linewidth of protein signals measured with different ligand concentrations. One of the most common applications of titration experiments consists in estimating dissociation constants (K_d_) (Fielding, 2003; Thordarson, 2011).

Analysis of 1D or 2D NMR titration experiments requires to process a large number of signals, a tedious, error prone process. In most studies, data processing and analysis are performed via independent (and mostly non released) scripts (Shortridge *et al*., 2008; Günther, 2002; Mittag *et al*., 2003), and only a few Matlab scripts are available for detailed analysis of particular titration experiments (Günther, 2002; Kovrigin, 2012; Waudby, 2016). There is thus the place for a generic and flexible tool to process and analyze NMR titration experiments, preferably implemented in Bruker BioSpin TopSpin™ – the main acquisition software used by the NMR community – to provide an end-to-end solution in a unified framework.

Here, we present Interact, a Python script accessible within TopSpin, which includes the following features:

- Automated peak picking and annotation of successive 1D or 2D spectra. Signals can be processed individually (basedon the spectral range of the currently displayed spectrum), or in batch (from a text file defining the spectral range of the signals of interest).
- Estimation of chemical shift(s), linewidth(s) and intensity of each signal in individual experiments, and calculation of the impact of the different ligand concentration on these parameters.
- Estimation of parameters informative of the protein-ligand interaction by integrating individual results. Thermodynamic (dissociation constant), structural (slope and angle of the ligand-induced changes of chemical shifts) and kinetic (ligand-induced changes of line width) information is extracted using optimization procedures, with visual inspection of fitted and experimental data.
- Sensitivity analyses are performed to determine the precision on the estimated parameters.
- Interact can be applied to all types of 1D and 2D NMR spectra and all nuclei.

## Methods and implementation

The general workflow implemented in Interact is shown in Figure 1, using 2D NMR titration experiments as illustrative example. Briefly, this workflow consists in i) peak picking and annotating each spectra, ii) processing the signal(s) of interest, and iii) integrating the results of the different experiments to infer thermodynamic, kinetic and structural information on the system under study. Several options can be provided by the user to easily adapt the workflow as well as the processing parameters to each situation. Report files gathering all the processing results are generated, thereby facilitating further analysis via existing and forthcoming alternative approaches (e.g. using tailored, case-specific protein-ligand interaction models). Additional details on the methods and on Interact usage can be found in the tutorial provided in Supplementary Data.

**Fig. 1.**
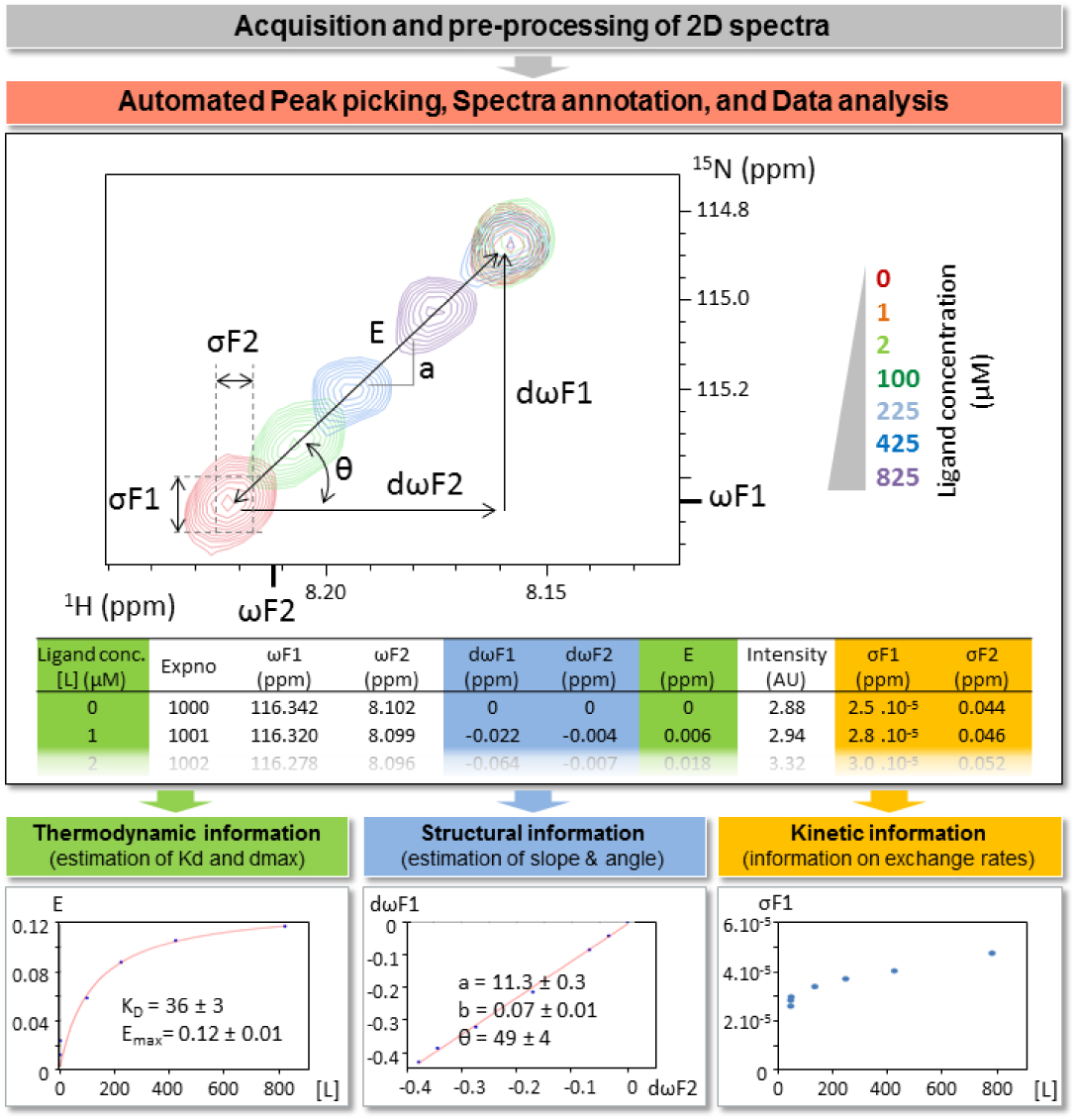
Workflow implemented in InterAct to analyze 2D NMRMR titration experiments. NMR spectra are first acquired, Fourier-transformed, phased, baseline-corrected and referenced using TopSpin. InterAct then performs peak picking and spectra annotation, and extracts several parameters of each signal of interest in each experiment *(ωF1, ωF2:* chemical shifts; *σF1, σF2:* linlinewidths; *I*: intensity). These results are then integrated to infer thermodynamic, structural and kinetic aspects information *(dσF1, dσF2, dωF1, dωF2:* ligand-induced changes of linewidths and chemical shifts; *E:* Euclidian distance; *a, θ:* slope and angle of changes of chemical shifts; *Kd:* dissociation constant).

### Input data

Titration experiments consist in analyzing samples prepared with a fixed protein concentration and different ligand concentrations. The resulting 1D or 2D NMR spectra must be Fourier-transformed, phased, baseline-corrected and referenced in TopSpin before running Interact.

### Signal analysis

Interact automatically detects the number of dimensions and extracts chemical shifts (*ωF1* and *ωF2* in Figure 1), linewidths (*σF1* and *σF2*), and intensity (*I*) for the signal(s) of interest in each experiment. Peak picking and spectral annotation are performed using TopSpin routines. For 1D spectra, the linewidth is also estimated using TopSpin. For 2D spectra, the linewidth is estimated in each dimension by fitting the experimental spectra to 2D Lorentzian or Gaussian (with or without rotation) models. Plots are generated for visual inspection of fitted and experimental spectra. Processing results are stored in tabulated text files.

### Data integration

Interact integrates the results obtained for each signal in the different experiments to extract additional information on the protein-ligand interaction. Changes of chemical shifts (*dωF1, dωF2*, and the corresponding Euclidian distance *E*, Fig.1) and linewidths (*dσF1* and *dσF2*) induced by the different ligand concentrations are measured in every spectrum. These parameters are exploited further to infer structural and thermodynamic information via fitting procedures. For each signal, the dissociation constant (*Kd*) is estimated using a two-state interaction model described by the following function (Williamson, 2013):

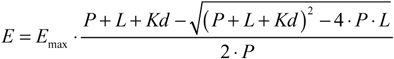

where *E* is the euclidean distance of the chemical shift caused by the ligand at concentration *L, E_max_* is the maximal euclidean distance (i.e. observed at ligand saturation), *P* is the protein concentration. Euclidean distance *E* is calculated as follow:

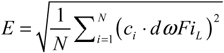

where *N* is the number of dimension, *dωFi_L_* is the change of chemical shift (in ppm) in dimension *Fi* induced by the ligand at concentration L, and *c_i_* is a weighting factor equal to *γ_X_/γ_H_* (where *γ_H_* is the proton’s gyromagnetic ratio and *γ_X_* is the gyromagnetic ratio of the nuclei observed in the corresponding dimension).

For 2D NMR spectra, the slope *(a)* and angle *(θ)* of ligand-induced changes of chemical shifts in each dimension are also estimated by linear regression.

For each fitting process, standard errors on the parameters are calculated from the estimated covariance matrix, and plots are generated for visual inspection of simulated and experimental data. Processing results are stored in tabulated text files.

### Implementation

This workflow has been implemented in Python programming language (http://python.org). Peak picking and annotation are performed by a TopSpin Python module, and data extraction, fitting and visualization are performed by external Python scripts. This enables seamless usage of Interact on Windows, MacOS and Linux, the three platforms supporting TopSpin. The source code is distributed under OpenSource license at http://github.com/MetaSys-LISBP/Interact.

## Example

Interact has been used to investigate the interaction between a short heparin oligosaccharide (degree of polymerization dp4) and the TauF4 fragment (Fauquant *et al*., 2011) to better understand how the poly-anion can drive the aggregation of Tau into fibers resembling the filaments that characterize patients with Alzheimer’s disease (Pérez *et al*., 1996). Briefly, 7 samples were prepared with a fixed concentration of TauF4 (100 μM) and different concentration of the dp4 heparin fragment (varied from 0 to 800 μM). 2D TROSY NMR spectra were acquired and pre-processed in TopSpin. Using Interact, the entire dataset was automatically processed – including annotation, peak picking and fitting of a total of 259 signals (37 signals on each of the 7 spectra), and data integration – in less than 2 hours on a personal computer.

Interact is thus capable of efficiently processing the large datasets that can now be acquired with modern, high-throughput NMR technologies.

By allowing end-to-end integration of all steps (from acquisition to spectra analysis) in a single software, Interact is expected to facilitate the analysis of existing and forthcoming 1D and 2D NMR titration experiments, thereby supporting fundamental advances in our mechanistic understanding of protein-ligand and protein-protein interactions.

## Acknowledgments

We thank Drs I. Huvent (Lille, France) and C. Wang (Troy, USA) for providing the Tau fragment and the heparin oligosaccharide, respectively, and Dr X. Hanoulle (Lille, France) and N. Cox (Toulouse, France) for testing the Interact software. MetaToul (Toulouse metabolomics & fluxomics facilities, www.metatoul.fr) is gratefully acknowledged.

## Author Contributions and Notes

G.L. and P.M. designed the study, P.M. developed the software G.L performed the experiments, P.M. and G.L. contributed to data and wrote the manuscript.

The authors declare no conflict of interest.

This article contains supporting information online.

